# Patterns of mortality in domesticated ruminants in Ethiopia

**DOI:** 10.1101/2022.07.04.498471

**Authors:** Giles T. Innocent, Ciara Vance, David Ewing, Iain J. McKendrick, Solomon Hailemariam, Veronica R. Nwankpa, Fiona K. Allan, Christian Schnier, Andrew R. Peters

**Author notes:** **Correspondence:** Andrew R. Peters.

## Abstract

**Background:** In Ethiopia, there are limited data on the causes of ruminant mortality and reproductive losses which is hampering the development of interventions to improve livestock health. The present study aimed to establish the causes and magnitude of mortality and reproductive losses in cattle, sheep and goats across all smallholder production systems in Ethiopia.

**Methods:** The study, consisting of three data sources, was conducted to collect data on mortality and reproductive losses in cattle, sheep and goats. Nine regional states; Afar, Amhara, Benishangul Gumuz, Gambela, Harari, Oromia, Southern Nations, Nationalities, and Peoples’ Region, Somali, and Tigray were surveyed, as well as the city administrations of Dire Dawa and Addis Ababa. Questionnaire surveys were conducted with farmers and key stakeholders, as well as data collected from regional diagnostic laboratories. Data from the Living Standards Measurement Study were also included. Generalised linear mixed and hurdle models were used for data analysis, with results summarized using predicted outcomes.

**Results:** Analyses indicated that most herds experienced zero mortality and reproductive losses, with rare occasions of larger losses. Diseases causing deaths varied greatly both geographically and over time. There was little agreement between the different datasets. While the models aid the understanding of patterns of mortality and reproductive losses, the degree of variation observed limited the predictive scope.

**Conclusions:** The models revealed some insight into why mortality rates are variable over time and are therefore less useful in measuring production or health status, and it is suggested that alternative measures of productivity, such as young stock mortality, would be more stable over time and likely more indicative.

## 1 Introduction

Livestock mortality is of great importance globally, and especially in low- and middle-income countries (LMICs) where livestock often have multiple roles. Livestock mortality has typically been considered one of the most important measures of population dynamics and productivity of livestock production systems (Putt et al., 1987), therefore high mortality risks can be considered a significant constraint in traditional livestock systems in sub-Saharan Africa (sSA) (Otte and Chilonda, 2002). Stock lost is a loss in nutrition, livelihood, and genetics, as well as lost wealth (Wong et al., 2021).

In Ethiopia, there is consistently low ruminant productivity and profitability (Shapiro et al., 2017). Between 2005 and 2015, average annual mortality in cattle and small ruminants was reported to be 7% and 20%, respectively (Central Statistical Agency, 2017a, 2017b). Infectious diseases are frequently reported in household surveys as being major constraints on smallholder productivity (Legese et al., 2014) and annual disease-associated losses are estimated at US $150 million (Berhanu, 2002). The rate of abortion in Ethiopian cattle is reported to range from 2.2-29%, varying across regions (Gizaw et al., 2007; Tesfaye and Shamble, 2013; Benti and Zewdie, 2014; Regassa and Ashebir, 2016; Siyoum et al., 2016), where an abortion rate of 2-5% per annum is considered to be expected (Tulu et al., 2018). In central Ethiopia, 57% of farmers have reported abortions in their flocks (Gebremedhin et al., 2013), with the reported mean annual abortion/stillbirth rate for Ethiopian flocks ranging from 3-36% over different ecosystems (Fentie, 2016; Gebretensay et al., 2019; Jones et al., 2020). Where the rate of abortion is between 2-5%, it is suggestive of underlying endemic disease, but when the rate exceeds this, it is important to investigate likely causes (Menzies, 2011). Despite this, and the reported high rates, the causes of abortion are largely unexplored (Teshale S et al., 2007; Esubalew et al., 2020). Overall, information on small ruminant abortions in Ethiopia is considered lacking (Gojam and Tulu, 2020).

Smallholder production systems in Ethiopia are broadly classified into three categories: mixed crop-livestock, pastoral and agopastoral systems. Mixed crop-livestock farming is the predominant system in Ethiopia, combining crop cultivation and livestock production. There is an emphasis on crop cultivation, with low numbers of livestock per household. For example, in Central Ethiopia, the average number of livestock per household is 4.5 cattle, 1.1 sheep and 0.5 goats (Duguma et al., 2012). Pastoral and agropastoral livestock production is the second most practiced system, predominantly in southern and eastern Ethiopia. There is no crop production in pastoral farming, and agropastoralism involves only limited crop and predominantly livestock production (Tegegne et al., 2013). Typically, per household, there are on average 21.1 cattle, 13.8 goats and 9.5 sheep (Tolera and Abebe, 2007).

Research and development of animal health projects and interventions in Ethiopia focus on transboundary or zoonotic diseases and those which affect trade, with relatively little attention given to endemic diseases, despite their negative effects on production, food security and livelihoods (Alemu et al., 2019). Indeed, there are limited quantitative data on the causes of ruminant losses in sSA which constrains the development and application of interventions to improve livestock health and productivity. The Centre for Supporting Evidence Based Interventions (SEBI), University of Edinburgh, was established to mobilise and apply evidence and data on livestock heath and productivity from LMICs. One of their initial programmes was to establish mortality rates and major causes of mortality in cattle and small ruminants, in Ethiopia, Nigeria and Tanzania. Outcomes have been published for studies carried out in Nigeria (Bolajoko et al., 2020) and Tanzania (Lankester, 2020). This study aimed to improve the available data for mortality and reproductive losses and their causes in cattle and small ruminants in states of Ethiopia, with the intention that appropriate interventions might be identified to reduce mortality and losses in cattle and small ruminants.

## 2 Materials and Methods

### 2.1 Ethics statement

No animals were involved in this study. Verbal informed consent was obtained from each participant prior to commencement of the survey. They were informed that their participation was voluntary, that their responses would be kept confidential and no names or identifying information linking them to the survey would be disclosed. In addition, they were assured that they were free to terminate their participation at any time. Surveys were read out to participants in the local language and responses translated into English by local translators.

At the outset of the study, ethical review was unintentionally omitted, however full details and supporting documentation of the study were submitted for retrospective ethical review on 21^st^ February 2022 by the Human Ethical Review Committee, University of Edinburgh, the outcome of which stated that had the study been reviewed prior to commencement, ethical approval would likely have been granted.

### 2.2 Datasets

A total of three datasets were used in the study: a Farmer Survey (FS), the Living Standards Measurement Study (LSMS) from the World Bank, and the Disease Outbreak and Vaccination Reporting (DOVAR) dataset. Each dataset contained data on number of deaths, state, and species. Species included were cattle, sheep and goats or ‘small ruminants’. Datasets also had varied additional information (Table 1), however, these were the only data that were consistent for all three datasets. The term ‘herd’ is used, hereafter, to refer to the varied reporting units, including ‘households’, ‘holdings’, ‘herds’ and ‘flocks’.

**Table 1.**
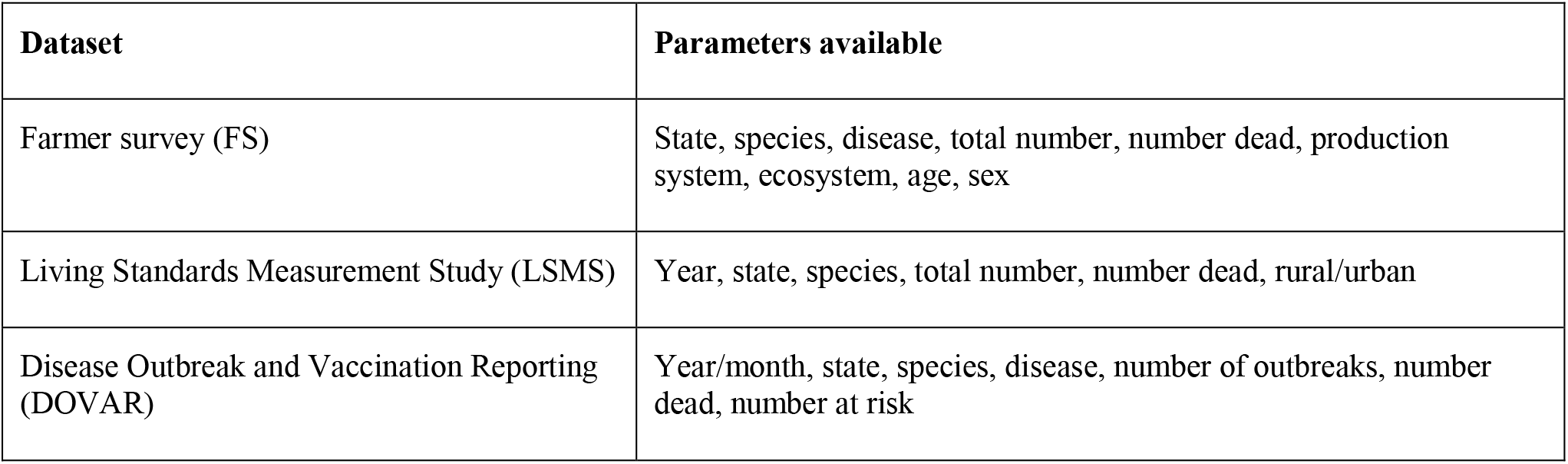
Data available in each dataset.

For numerical reasons, we aggregated sheep and goats in a single small ruminant (SR) class. Due to data availability, we were only able to analyse abortions in the FS.

Data were sparse and there was a lack of independence of potential explanatory variables, such as Age, Sex, Disease, State, Ecosystem and Production System. Statistical models did not converge when ‘Disease’ was included as a variable, therefore disease variables were recoded and grouped as shown in Table 2.

**Table 2.**
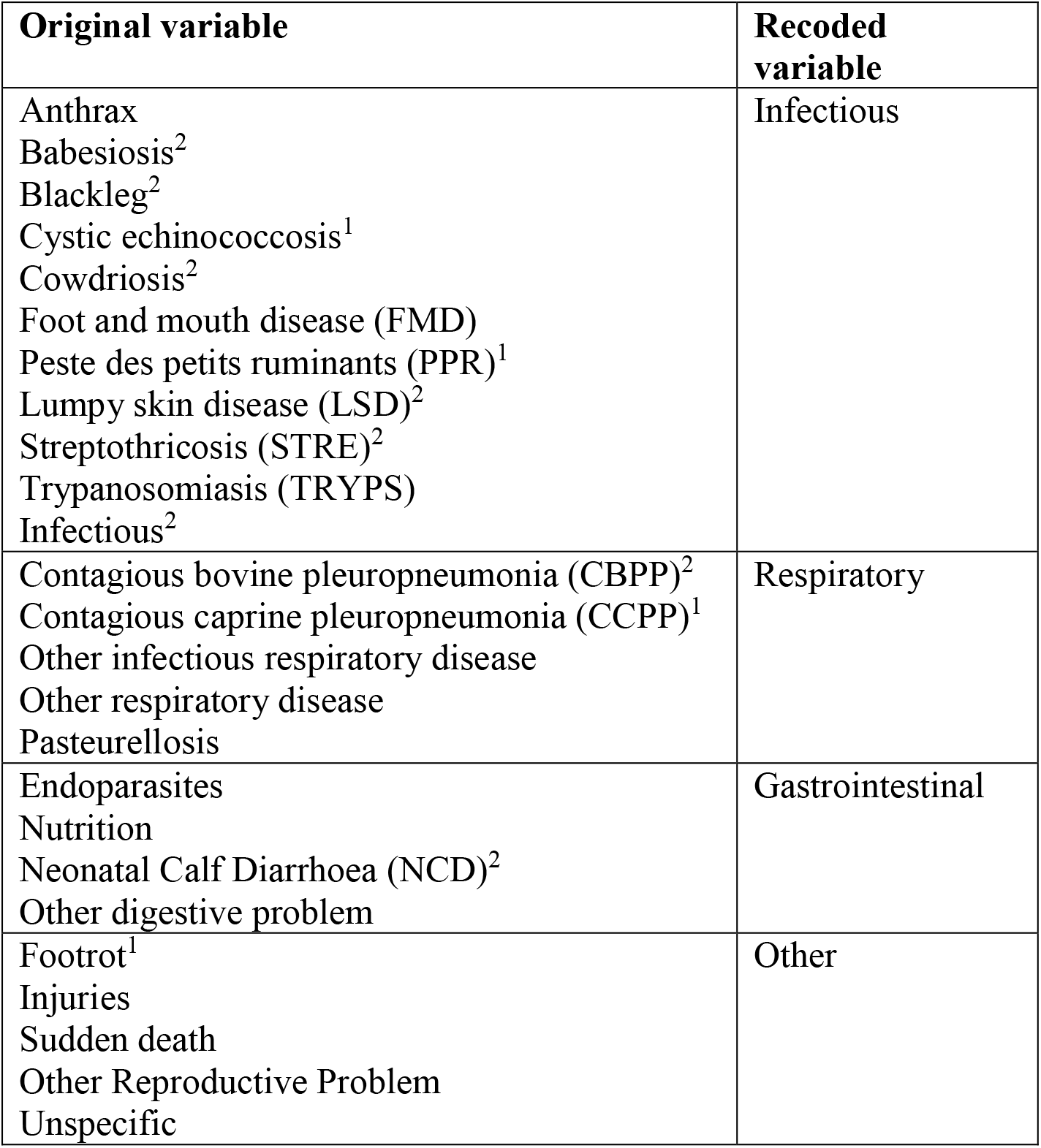
Grouping of disease variables. ^1^Disease only recorded in small ruminants. ^2^Disease only recorded in cattle.

#### 2.2.1 Farmer surveys

Farmers were recruited in purposively selected zones, districts and wards in all nine regional states^1^ and in Dire Dawa city administration, based on livestock population, and included different production systems and agro-ecological zones (AEZ). Two peasant associations (PA) were additionally purposively selected, based on accessibility and ruminant population. Ten households/herds per PA were selected for interview and group discussion, based on their experience in the livestock sector. Households or producers (farmers, pastoralists and urban producers) were selected primarily based on their availability and accessibility to the study teams. A total of 382 herds were sampled.

Structured questionnaire surveys (Supplementary Material) were designed by SEBI and conducted with farmers between October and December 2017. Five study teams conducted the surveys with farmers and pastoralists, collecting information on demographics, causes of mortality and reproductive performance losses in cattle and small ruminants. Semi-structured questionnaires were carried out with key informants and stakeholders. Primary information was collected from producers, via participatory appraisal tools, focus group discussions, key informant interviews and visual observations. Secondary information (published and unpublished documents) was collected from livestock district offices, regional laboratories, universities, ranches, and private and government farms.

Herd data collection categorized cattle and small ruminants by age and sex. Ages were classified as young stock (≤2 years), juveniles (>2-3 years) and adults (4+ years). Information was captured for the previous year (2016) on reproductive performance losses was collected from farmers, pastoralists, artificial insemination centres, and livestock offices through discussion. The incidence of stillbirths was not raised and not specifically included; producers rather reported pregnancy terminations as abortions. Information on causes of mortality in cattle and small ruminants were also collected from district offices, regional livestock departments and regional veterinary laboratories for the preceding year.

Verbal consent was gained from each participant prior to commencement of the survey; all participants chose to participate. The survey was explained verbally in the local regional language, and verbal consent was obtained.

#### 2.2.2 Living Standards Measurement Study (LSMS)

The LSMS-Integrated Surveys on Agriculture (LSMS-ISA) are household survey projects that collaborate with the national statistics offices in partner countries, of which Ethiopia is one. The LSMS-ISA project supports the design and implementation of the Ethiopia Socioeconomic Survey and is implemented every two years (The World Bank, 2021).

For our study, we used data collected from rural households (n = 4,000) in 2011/2012 (wave 1) and the same households revisited and expanded to include 1,500 urban households (n = 5,500) in 2013/2014 (wave 2). Sampling followed a stratified random sampling scheme intended to select households according to population size. For information on sampling design, refer to The World Bank (2021).

Data were collected face to face via a livestock questionnaire interview, administered where at least one member of the household was involved in rearing livestock. Information was collected for the previous 12 months about animal holdings and costs, and production, cost and sales of livestock byproducts. Data were collected for deaths but not abortions. Completed questionnaires were sent to the Central Statistical Agency in Addis Ababa. Questionnaire design for both waves of data collection and details for field work implementation are described previously (CSA & LSMS World Bank, 2017; The World Bank, 2021).

#### 2.2.3 Disease Outbreak and Vaccination Reporting (DOVAR)

DOVAR collated data from seven regional diagnostic laboratories within Ethiopia (Mekele, Asosa, Bahardar, National Animal Health Diagnostic and Investigation Centre (NAHDIC) in Sebeta, Shola, Dire Dawa, and Soddo) as well as from the Department of Disease Outbreak Investigation and Vaccination in Addis Ababa.

The data were collected by the laboratories monthly between 2009 and 2015, and annually in 2016 and 2017. The laboratories collected samples from sick animals during outbreaks for analysis as well as conducting post-mortems. Data in our study were taken from the laboratory registers, between December 2017 and early 2018.

### 2.3 Data analysis

Initially, where the total number of animals was recorded, a binomial model was proposed for these data, that is, the number of deaths or abortions relative to the numbers of animal kept. However, a large number of records indicated more deaths or abortions than animals kept. This uncertainty about numbers may represent where a number of animals were kept at the beginning of the year, some animals died, and the remaining animals are reported as the number kept. Therefore the binomial model approach was abandoned, in favour of using the number of animals as a covariate in the model as a more robust approach. This choice of model means, however, that even where the number of animals per herd is specified, it is possible for the model to predict that a greater number of deaths will occur.

Therefore, all datasets were analysed using a generalised linear mixed model, (GLMM), specifically a Poisson model with overdispersion fitted as a random intercept associated with herd and zero-inflation, if it improved the model fit. Use of a mixed model allowed for the variation between herds and the random effect represents the accumulation of many small herd-level effects that were not measured, such as training, cleanliness etc. Where possible, an offset term was included to account for the differing numbers of animals in each herd. Where herd size was not available either a Poisson or Negative Binomial model was used, whichever had the better fit.

In practice, an initial model was proposed for each dataset, with a Poisson error structure with a random effect based on herd ID, where this was available. The model was compared to a similar model allowing for an increased number of zero observations using glmmTMB (Brooks et al., 2017) in R (R Core Team, 2019), and where there was support for the latter model, an initial logistic regression model was considered, using glm, and the optimum model for zero-non-zero observations, including explanatory variables, was identified. A conditional (non-zero observation) model was then fitted using glmmTMB. The non-zero observations were modelled as a zero-truncated Poisson distribution with herd fitted as a random intercept, and the optimum set of explanatory variables identified. Formally this can be described as a “hurdle” model, reflecting the two stages of the fitting process.

Models containing various combinations of explanatory variables were compared using Akaike’s Information Criterion (AIC), with the model with the lowest AIC selected as the best fitting model, subject to model assumptions being satisfied. Explanatory variables in the final model are presented on a case-by-case basis as the variables available for inclusion were not consistent across species and outcomes. Where models would not converge, a one-way non-parametric test equivalent to a parametric analysis of variance, a Kruskal-Wallis test, was used. Model results were summarised by predicted values based on the best fitting model and presented as box and whisker plots.

## 3 Results

### 3.1 Farmer survey

The study involved 10,291 cattle (1,266 cross breeds) and 16,587 small ruminants (sheep and goats).

#### 3.1.1 Deaths in cattle

For cattle deaths (per annum), the final model selected was: the probability of observing zero or non-zero depended on Production System, State, Disease, Ecosystem and interactions between Production System and Disease, State and Disease, and Disease and Ecosystem. Taking this into account, the differences between non-zero counts could be accounted for by Production System, Ecosystem, Disease, Age of animal and interactions between Ecosystem and Disease, Disease and Age, and Ecosystem and Age. The complex relationships between State, Disease and Age group are shown in Figure 1, and State, Disease and Production System are shown in Figure 2.

**Figure 1.**
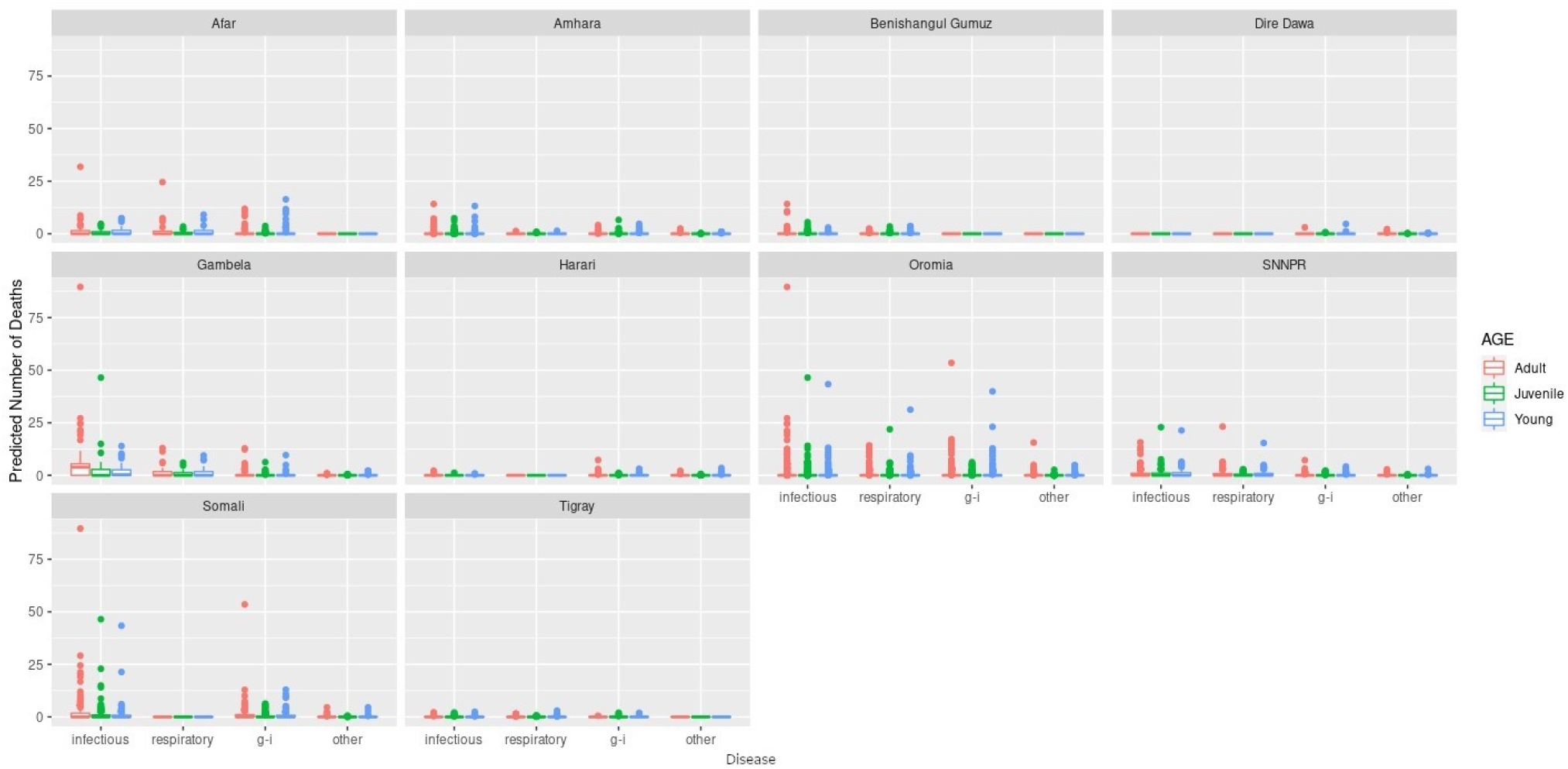
Predicted number of cattle deaths from relationships between State, Disease and Age group. Number of deaths is shown in the y-axis; State is shown in the top of each box; Age is represented by colour, where adults (4+ years) are shown in pink, juvenile (>2-3 years) are shown in green, and young stock (≤2 years) are shown in blue; and Diseases (recoded) are shown along the x-axis.

**Figure 2.**
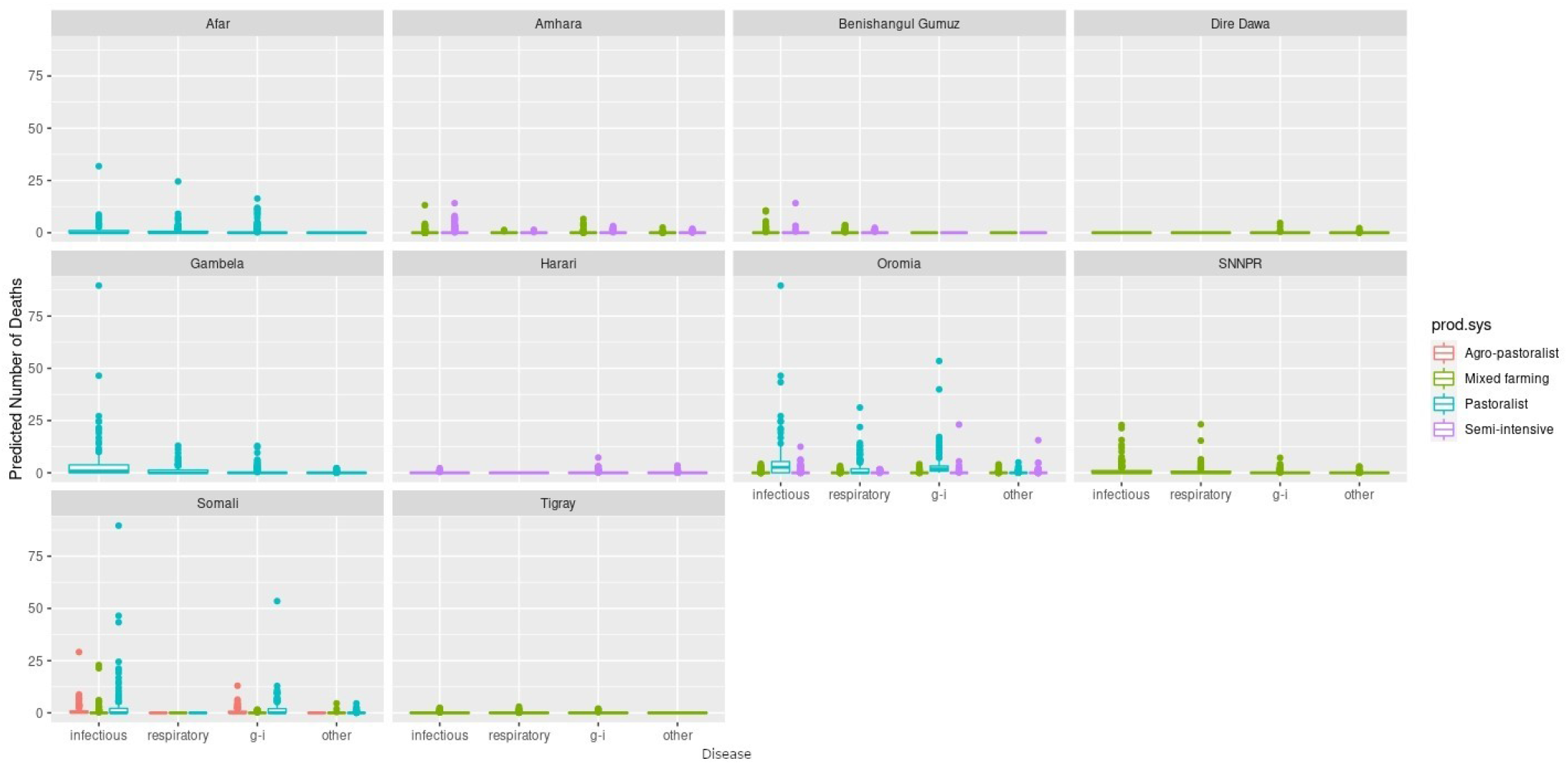
Predicted number of cattle deaths from relationship between State, Disease and Production System. Number of deaths is shown in the y-axis; State is shown in the top of each box; Production System is represented by colour as detailed in the legend; and Diseases (recoded) are shown along the x-axis.

Of all cattle herds included in the survey, 29% (109 of 382) had no deaths reported for any of the diseases. Generally, a greater number of deaths was predicted for older age groups, the states of Oromia and Somali, infectious diseases and pastoralist systems (Figures 1 and 2). Indeed, for a few pastoralist herds in Somali, Gambela and Oromia, the model predicted up to 90 deaths in adult cattle and up to 50 deaths in younger cattle due to infectious diseases.

#### 3.1.2 Abortions in cattle

There were a total of 3,064 records, of which 95% (2,897) were zero. The final models selected were: the probability of observing zero or non-zero events depended on State, Disease, Ecosystem, and interactions between State and Disease, Disease and Ecosystem, and State and Ecosystem; the count of non-zero entries depended on State, Production System, Disease, Ecosystem and an interaction between Disease and Ecosystem.

The results are presented as predictions. Figure 3 demonstrates how abortions vary by state, ecosystem and disease.

**Figure 3.**
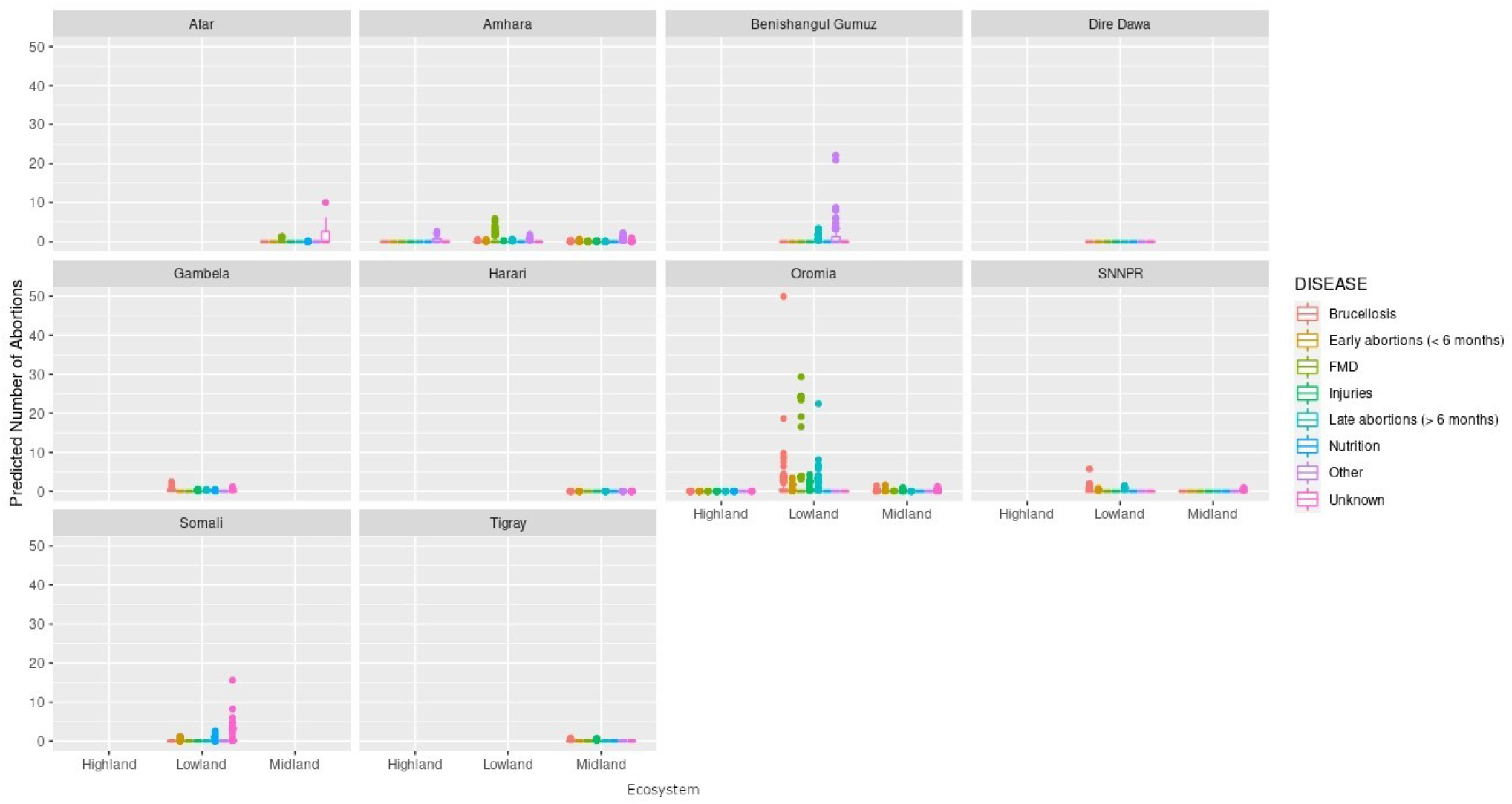
Predicted mean number of cattle abortions given State, Ecosystem and Disease. Number of abortions is shown in the y-axis; State is shown in the top of each box; Disease is represented by colour as detailed in the legend; and Ecosystems are shown along the x-axis.

The great majority of herds had no abortions predicted due to any disease in any of the states in any ecosystem. The highest number of abortions predicted for a herd were caused by Brucellosis in Oromia and mainly in the lowland regions, followed by FMD and late abortions (> 6 months) in the same state and ecosystem.

#### 3.1.2 Deaths in small ruminants

We included 16,587 small ruminants in the analysis, of which 866 (5%) died (57 of the 303 herds reported no deaths). Similarly to cattle, SR mortality was not uniformly distributed across states. Variables that were significantly related to mortality included Disease, State, Sex, Age, Total Number of animals, Production System and Ecosystem with several interactions between variables. As the fitted models included the effect of Total Number of animals held, this term was set to a standardized value of 10, close to the median value, to standardize predictions. Figure 4 demonstrates how deaths vary by State, Ecosystem and Disease, and Figure 5 demonstrates variation by State, Disease and Production System.

**Figure 4.**
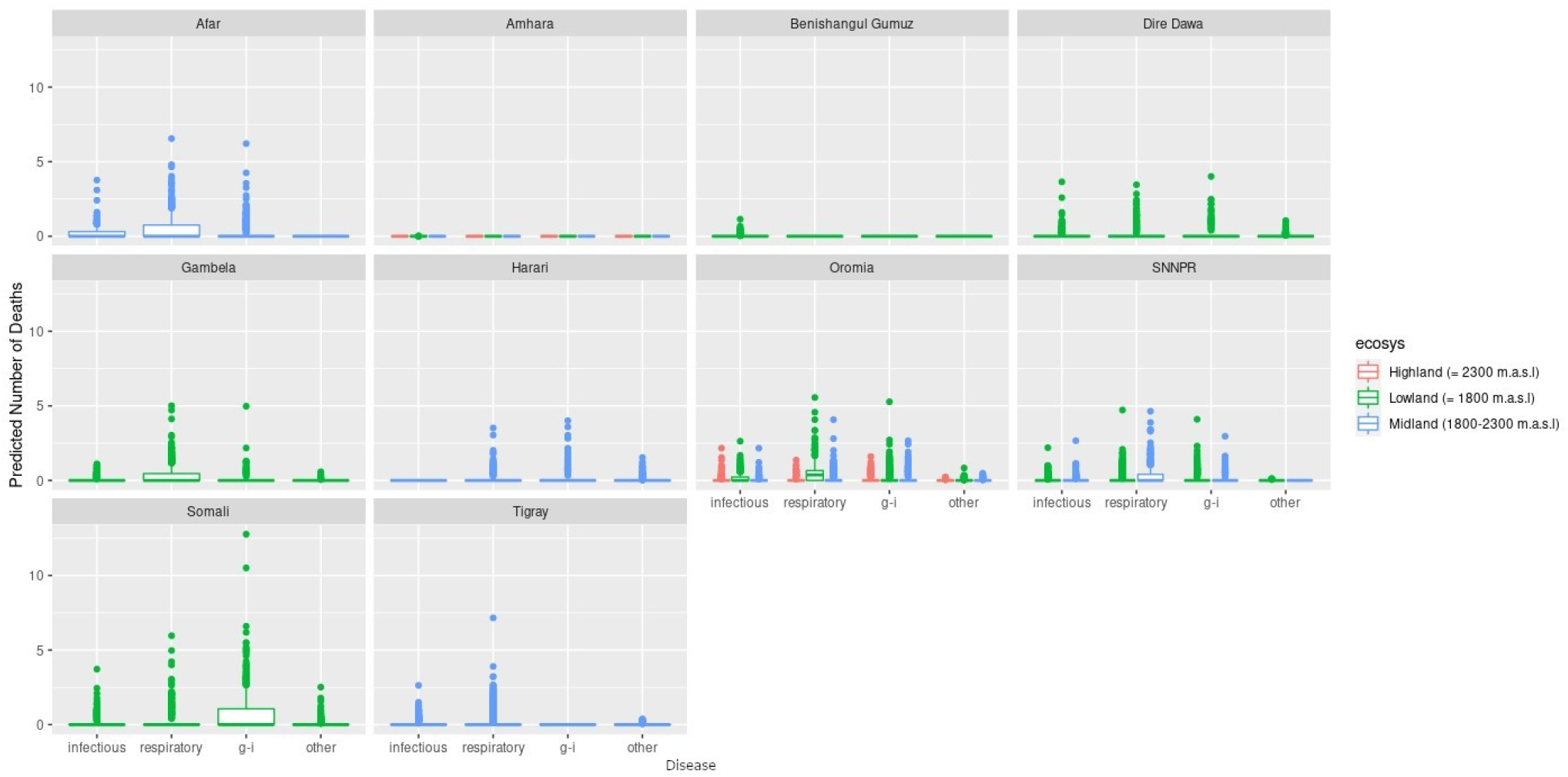
Predicted mean number of small ruminant deaths given State, Ecosystem and Disease. Number of deaths is shown in the y-axis; State is shown in the top of each box; Ecosystem is represented by colour as detailed in the legend; and Diseases (recoded) are shown along the x-axis. Results are presented as predictions across all herds, estimating standardized effects as if all herds had ten animals. m.a.s.l. = metres above sea level.

**Figure 5.**
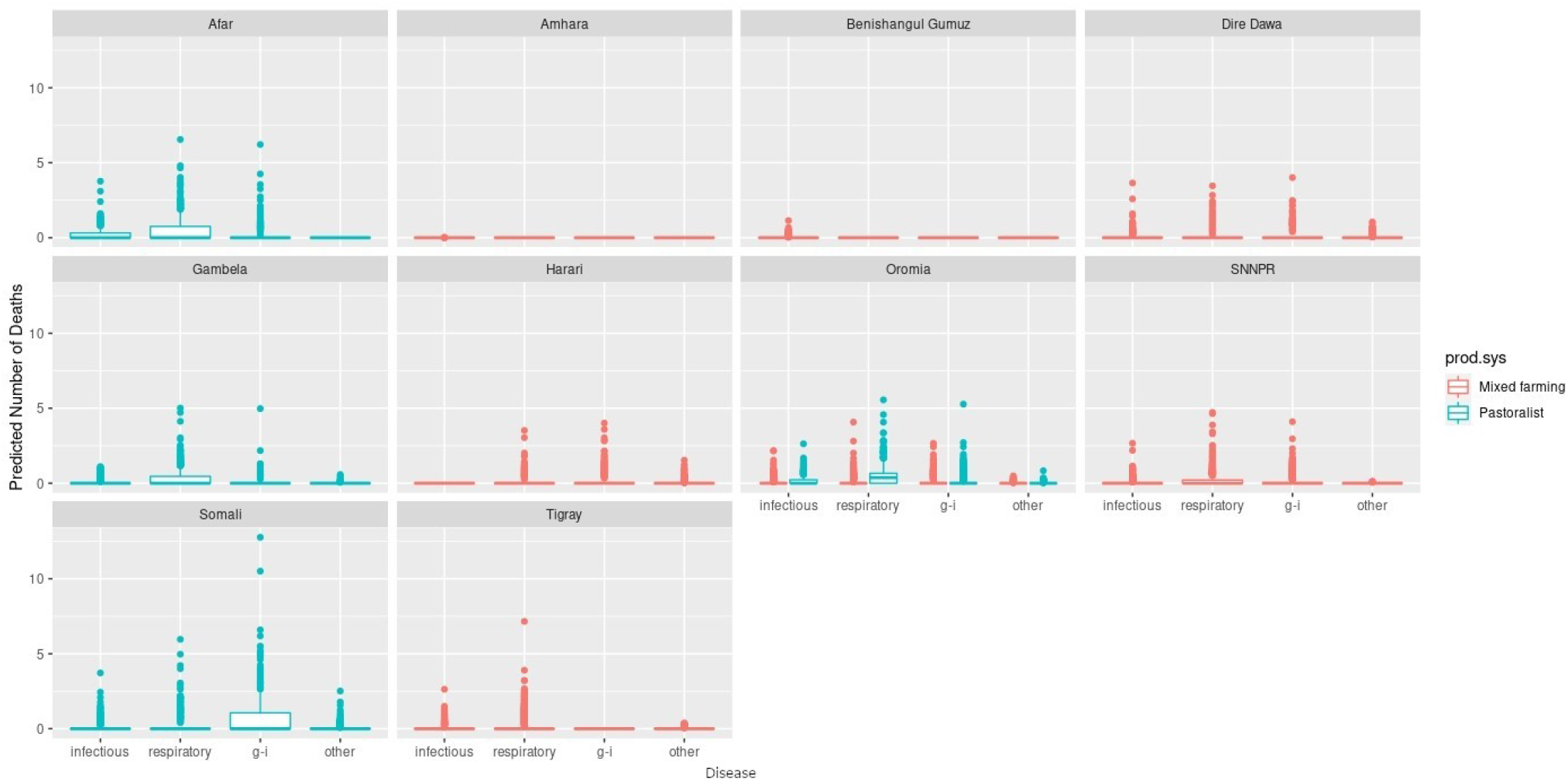
Predicted mean number of small ruminant deaths given State, Production System and Disease. Number of deaths is shown in the y-axis; State is shown in the top of each box; Production System is represented by colour as detailed in the legend; and Diseases (recoded) are shown along the x-axis. Results are presented as predictions across all herds, estimating standardized effects as if all herds had ten animals.

Most SR herds included in the survey were predicted to have no deaths due to any of the diseases. Increased mean number of deaths were associated with respiratory diseases, the states of Afar, Oromia and Somali, and pastoralist farming (Figures 4 and 5). However, in Somali, increased mean number of deaths in pastoralist herds were associated with gastrointestinal diseases.

#### 3.1.3 Abortions in small ruminants

Of 2,128 small ruminants, 104 (4.8%) had an abortion. The best-fitting model for zero against non-zero observations included Disease, Ecosystem, State, Total animals, and interactions between Disease and State, State and Total animals, Ecosystem and State, and between Disease and Total animals. The best-fitting model for number of cases, conditional on there being greater than zero cases, included Disease, State, Ecosystem, Total number of animals, and Production System, with no interactions. Results are presented as mean predictions across all herds (Figures 6 and 7).

**Figure 6.**
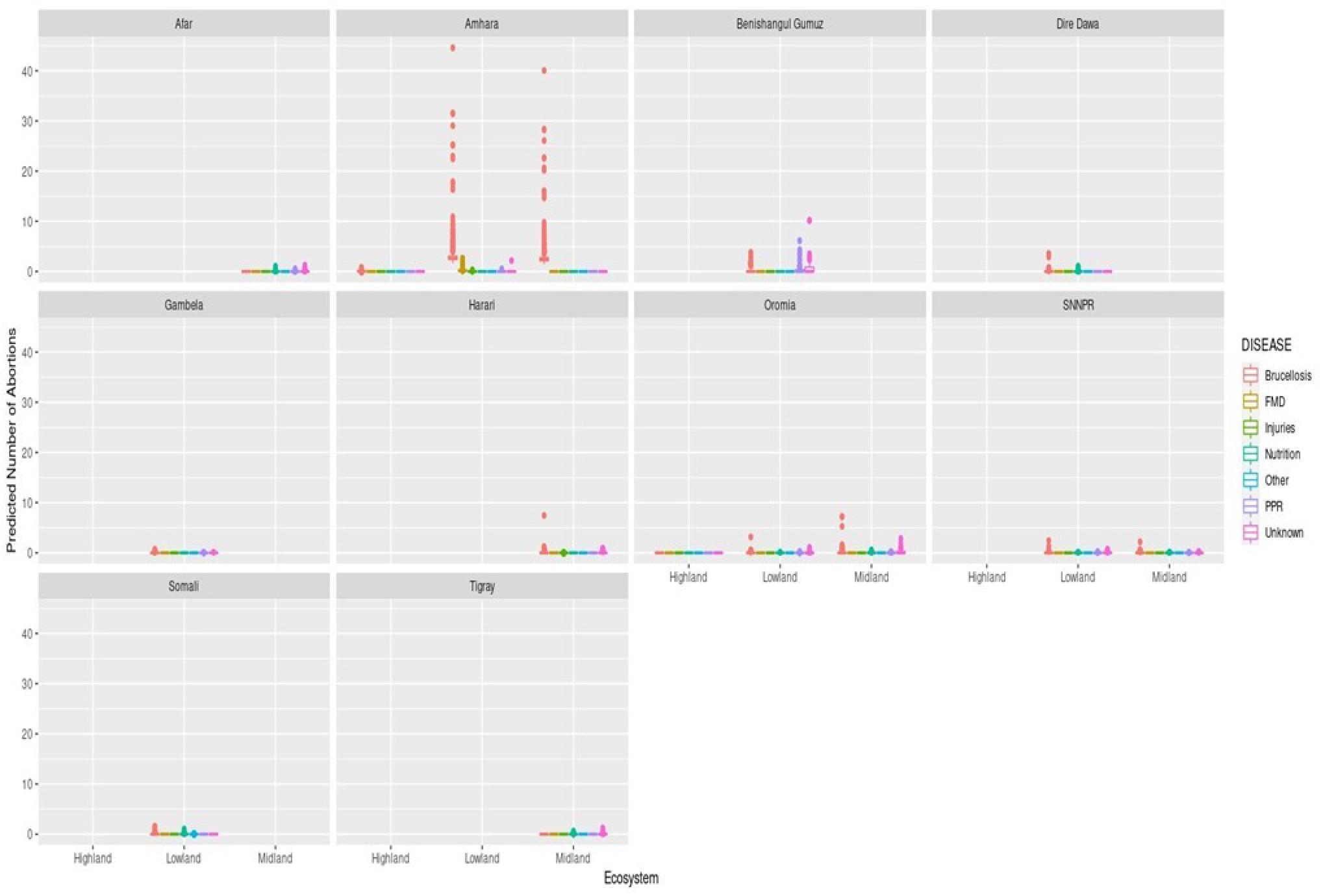
Predicted mean number of small ruminant abortions given State, Disease and Ecosystem. Number of abortions is shown in the y-axis; State is shown in the top of each box; Diseases are represented by colour as detailed in the legend; and Ecosystems are shown along the x-axis.

**Figure 7.**
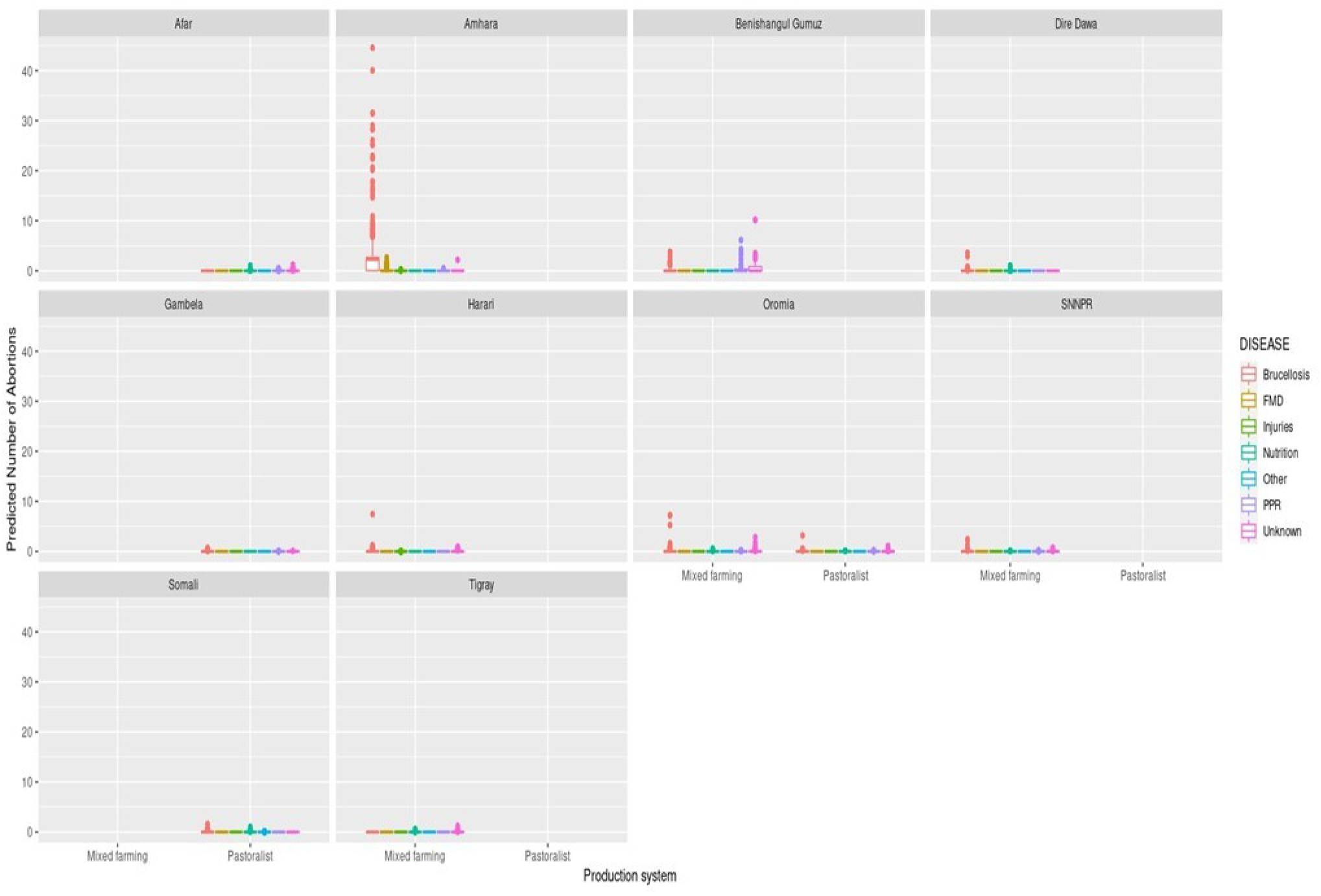
Predicted mean number of small ruminant abortions given State, Disease and Production System. Number of abortions is shown in the y-axis; State is shown in the top of each box; Diseases are represented by colour as detailed in the legend; and Production Systems are shown along the x-axis.

Most herds had no abortions due to any disease, in all states, ecosystems and production systems. In Amhara, herds situated in lowland and midland ecosystems were more likely to have abortions than herds in highland areas. Mean numbers of abortions due to brucellosis were predicted to be very high (>45) in lowlands and midlands, and mixed farming in Amhara. In contrast to the analysis of deaths, in Oromia there were marginally more abortions predicted in mixed farming than pastoralist systems.

### 3.2 Living Standards Measurement Study (LSMS)

#### 3.2.1 Cattle deaths

In this dataset there was a total of 11,650 non-duplicated records for 2011 and 2013. Of these, 7,081 records were deemed valid animal holdings, of which 6,182 recorded zero deaths (87%). Cause of death (disease) was not recorded.

The best-fitting model for zero versus non-zero observations included State, Year, Longitude, Elevation, Agroecology, Sex, and interactions between Year and Longitude, State and Longitude, State and Elevation, Longitude and Agroecology and between State and Agroecology as terms. The best-fitting model for the non-zero counts of deaths included Latitude and Year. Results are presented as mean predictions across all herds, standardizing the effects as if all herds had ten animals (Figure 8 and 9).

**Figure 8.**
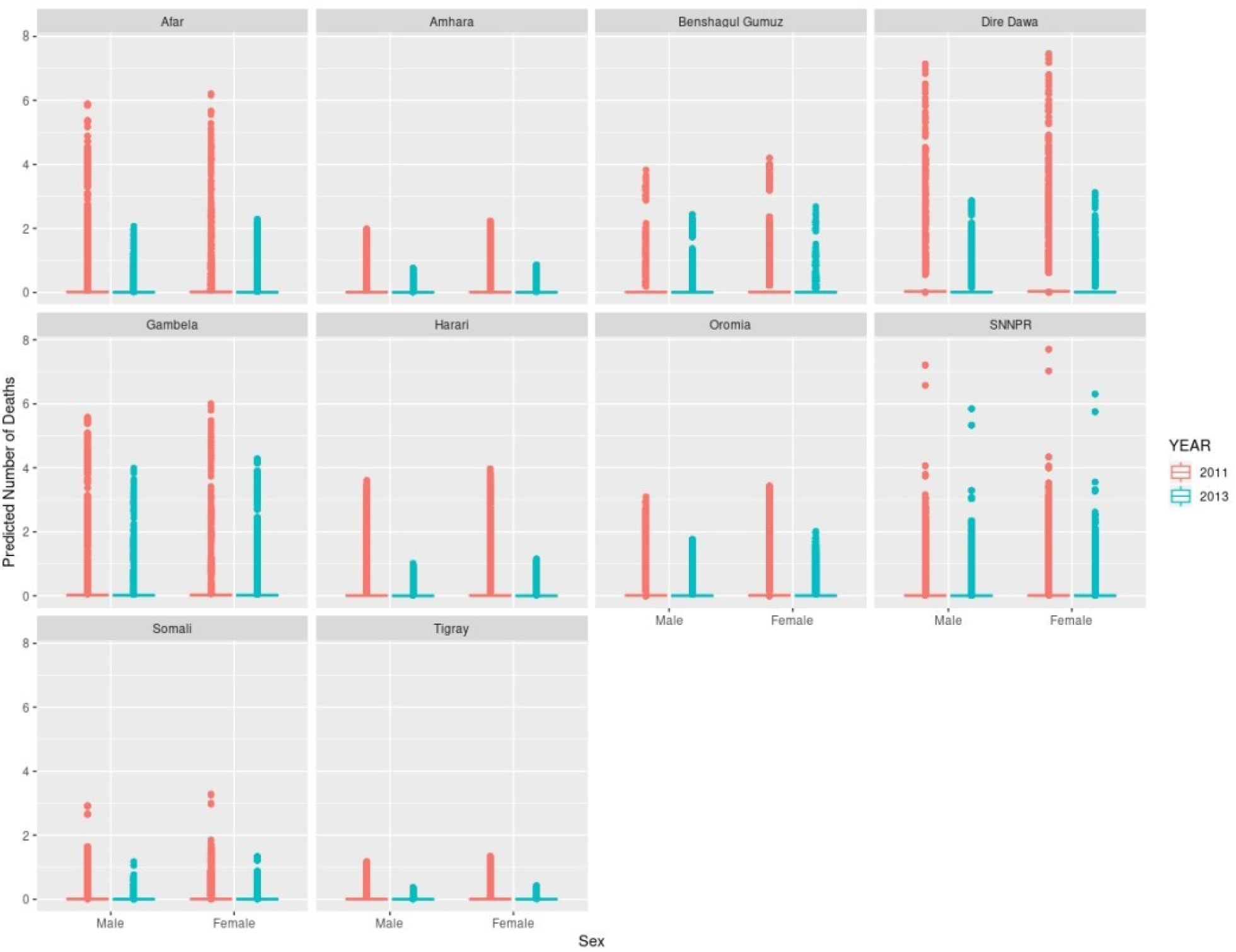
Predicted mean number of cattle deaths given State, Year and Sex. Number of deaths is shown in the y-axis; State is shown in the top of each box; Year is represented by colour, where 2011 is shown in pink and 2013 is shown in blue; and cattle Sex is shown along the x-axis. Results are presented as predictions across all herds, estimating standardized effects as if all herds had ten animals.

**Figure 9.**
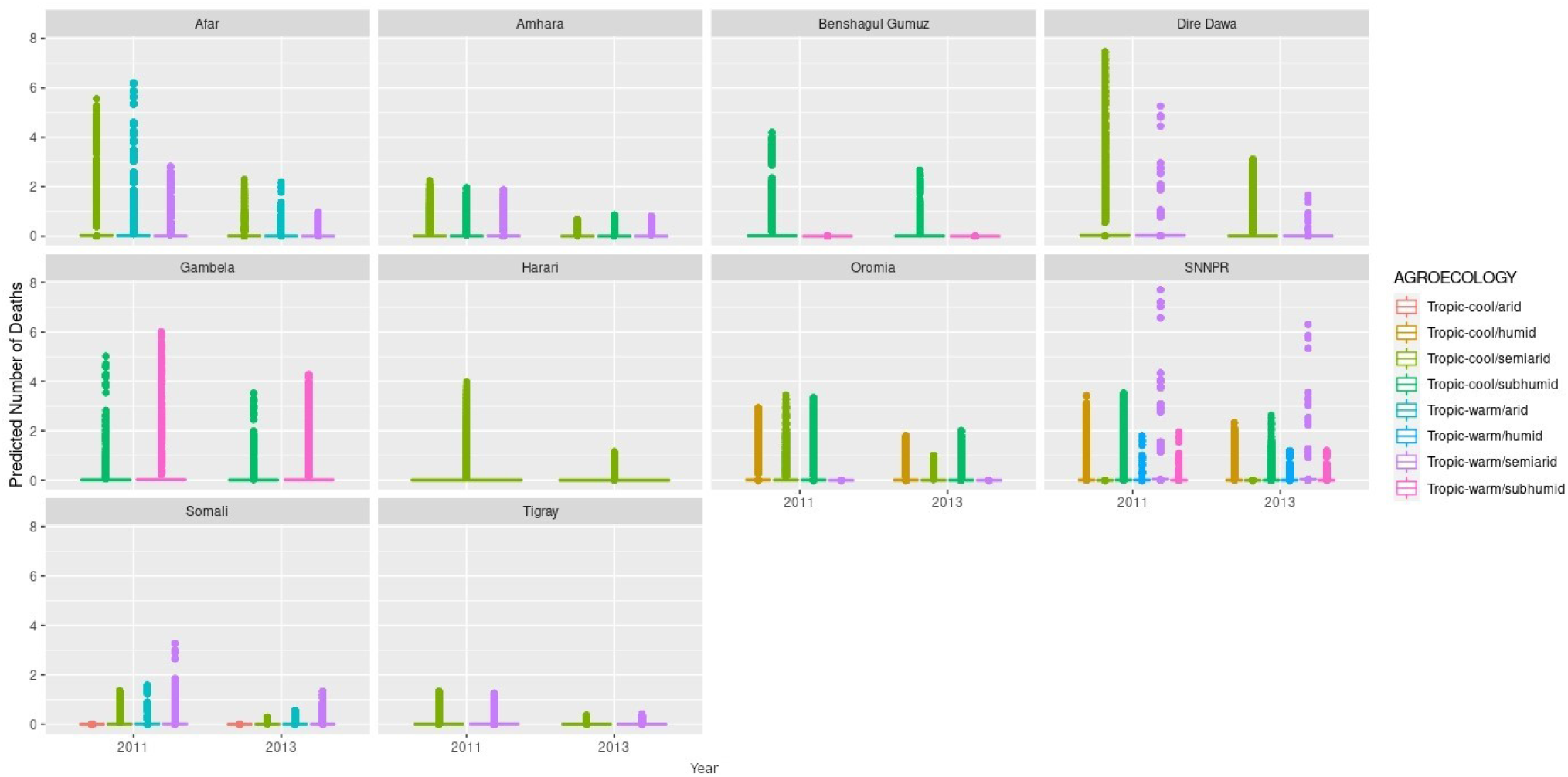
Predicted mean number of cattle deaths given State, Agroecology and Year. Number of abortions is shown in the y-axis; State is shown in the top of each box; Agroecology is represented by colour, as detailed in the legend; and Year is shown along the x-axis. Results are presented as predictions across all herds, estimating standardized effects as if all herds had ten animals.

In the model predictions (for a herd of 10 cattle), the majority of herds had zero deaths for both years, in all states and in both sexes. Where there were deaths predicted, there are major differences between states and years, and the effect of year varies between states. Dire Dawa had the highest mean number of cattle deaths predicted in 2011, for both male and female cattle, whereas Gambela had the highest number in 2013, again for both sexes. The lowest number of predicted deaths, for both sexes, and in both years, were in Tigray.

The SNNPR recorded data for six of the eight agroecological zones, the most of any state. Harari only recorded data for a single zone. The agroecological zone with the greatest risk of animals dying in 2011 is ‘tropic-warm/semiarid’ in SNNPR, followed by ‘tropic-cool/semiarid’ in Dire Dawa, whereas in 2013, it is ‘tropic-warm/semiarid’ in SNNPR again, followed by ‘tropic-warm/subhumid’ in Gambela. Overall, there were more predicted deaths in 2011 compared to 2013, for all states.

#### 3.2.2 Small ruminant deaths

There was a total of 23,250 non-duplicated records for 2011 and 2013. Of these, 6,457 records were deemed valid animal holdings, of which 5,305 recorded zero deaths (82%). As for cattle, cause of death (disease) was not recorded.

The best-fitting model for zero versus non-zero entries included State, year, Sex, Longitude, Agroecology, Elevation, and interactions between State and Elevation, State and Longitude, State and Agroecology, Longitude and Agroecology, Elevation and Agroecology, Year and Longitude, Elevation and Year, Elevation and Longitude, between State, Elevation and Agroecology and between State, Longitude and Agroecology as terms. This model therefore included six direct effects, eight two-way interactions and two three-way interactions.

The best-fitting model for non-zero counts included Longitude, animal Sex, Agroecology, Year and an interaction between Longitude and Year as terms, with a random effect of herd, and an offset of the natural logarithm of the adjusted number of animals held *(n* + 0.5 to account for zeros). Results are presented as predictions across all herds, estimating the effects if all herds had ten animals (Figure 10).

**Figure 10.**
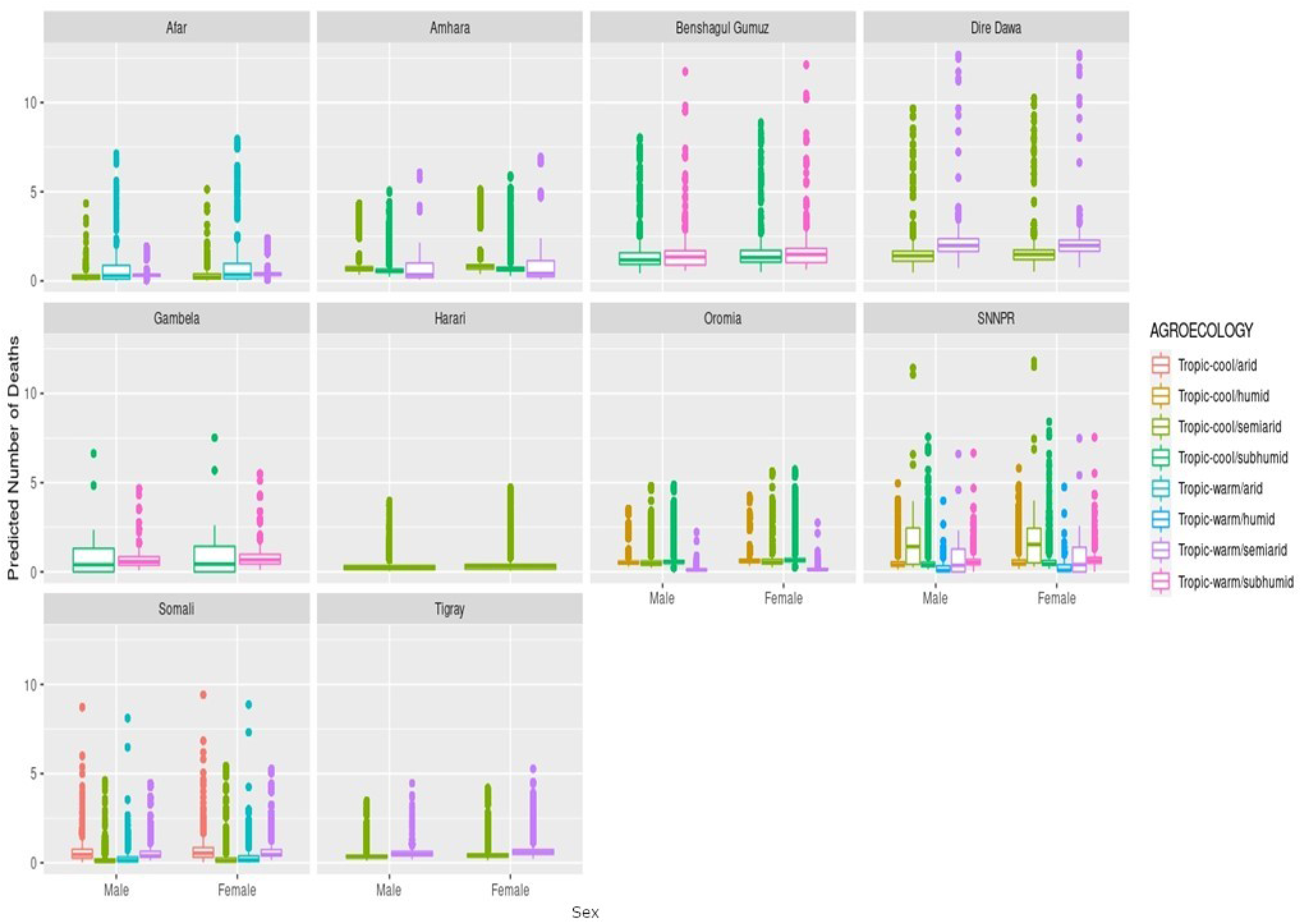
Predicted mean number of small ruminant deaths given State, Agroecology and Sex. Number of deaths is shown in the y-axis; State is shown in the top of each box; Agroecology is represented by colour, as detailed in the legend; and Sex is shown along the x-axis. Results are presented as predictions across all herds, standardising the effects if all herds had ten animals.

Median mortality in small ruminants for a herd of 10 animals was predicted to be under 3 for herds in all agro-ecological areas within the different states. Highest predicted mortalities indicated that complete herds perished. Increased mortality was seen in Benshagul Gumuz, Dire Dawa and Gambela, and female small ruminants were at slightly higher risk than males. There was no clear trend in the association of mortality with either humidity or temperature.

### 3.3 Disease Outbreak and Vaccination Reporting (DOVAR)

#### 3.3.1 Deaths in cattle

These data represent monthly returns of number of outbreaks, cases and deaths from notifiable diseases, at state level. If there were two outbreaks ongoing, it was possible to have two separate records for a single disease in a state in the same month, therefore data were aggregated by State, Disease and Year in order to be more in line with the other datasets analyzed, with values for all States, Diseases and Years fully calculated or inferred i.e. where no entry occurred in the data for a specific State/Disease/Year combination a zero was recorded, since no deaths were reported. A total of 1,386 records resulted, of which 1,060 were zero, 79 were >100 and 5 were >1,000 (maximum value 2,471).

The best-fitting model for whether any deaths were recorded or not depended on Year, State and Disease. Conditional models, that is models based using zero-truncated distributions on the non-zero data, failed to converge, so a Kruskal-Wallis test was used. Differences, statistically significant at the 5% level, were found between States (p <0.001), and also between Diseases (p <0.001), but not between years (p = 0.15). These differences are shown in Figure 11, where number of deaths is shown on the vertical axis on a log scale, as is appropriate for such skewed data.

**Figure 11.**
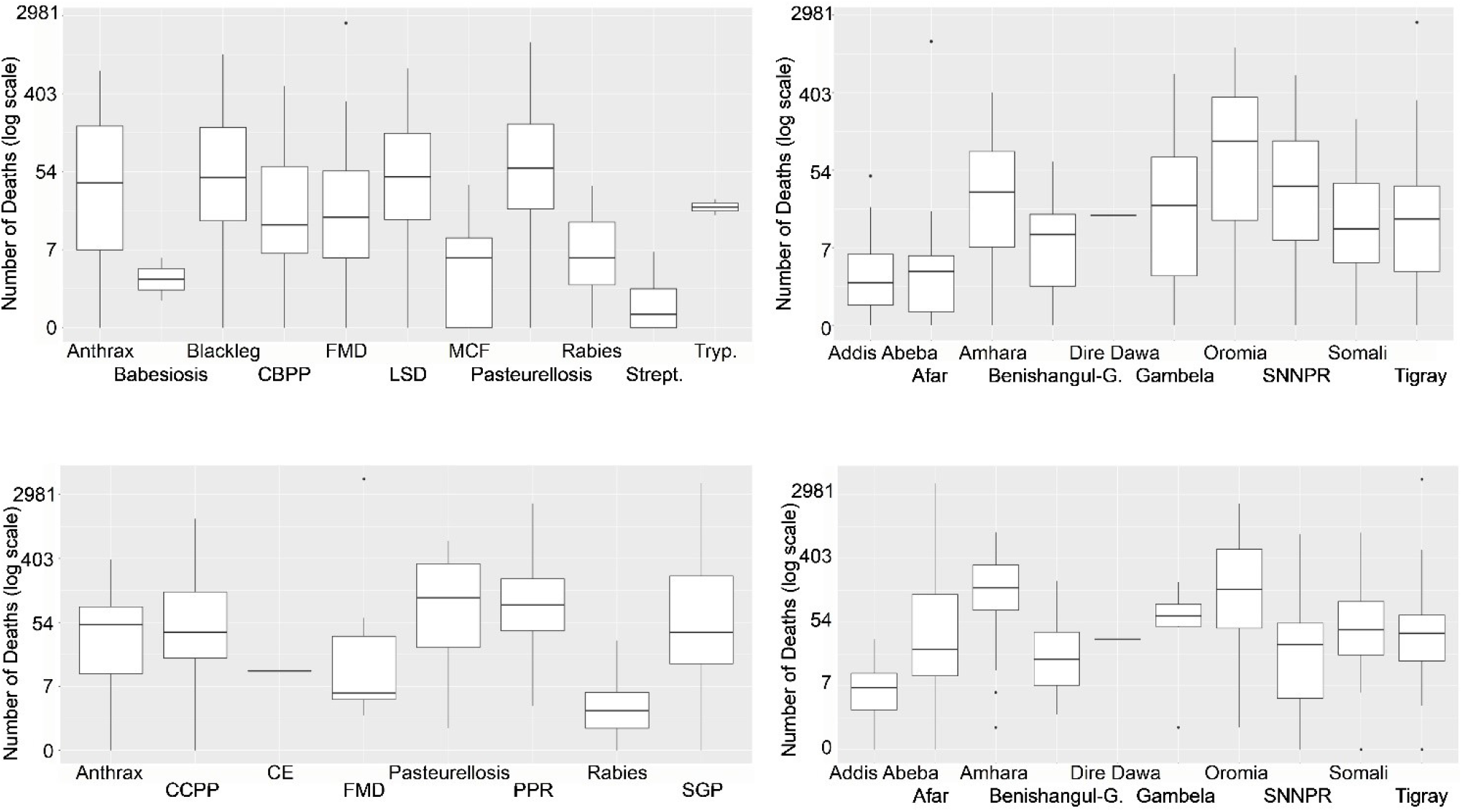
(**A**) Number of cattle deaths in outbreaks by Disease between 2009 and 2017, where deaths were reported. (**B**) Number of cattle deaths in outbreaks by State between 2009 and 2017, where deaths were reported. (**C**) Number of small ruminant deaths in outbreaks by Disease between 2009 and 2017, where deaths were reported. (**D**) Number of small ruminant deaths in outbreaks by State between 2009 and 2017, where deaths were reported.

#### 3.3.2 Deaths in small ruminants

As described above for the cattle data, these data were aggregated by State, Disease and Year. A total of 900 records resulted, of which 677 were zero, 80 were >100 and 6 were >1,000 (maximum value 4,838).

The best fitting model for whether any deaths were recorded or not depended on Year, State and Disease. Again, conditional models failed to converge, so a Kruskal-Wallis test was used. Differences, statistically significant at the 5% level, were found between States (p <0.001), and also between Diseases (p <0.001), but not between Years (p = 0.28). These differences are shown in Figure 11, where non-zero number of deaths is shown on the vertical axis on a log scale, as is appropriate for Poisson or negative Binomial data.

We can interpret this analysis as indicating that whether or not an outbreak occurs varies depending on the particular combination of State, Year and Disease, but should an outbreak of a specific disease be recorded, the mean number of small ruminants to die will be relatively consistent (independent of state, year or of disease).

For both cattle and SR, moderate variability was predicted for the number of deaths caused by different diseases and in different states, with a median of 7 to 55 cattle deaths and 7 to 403 SR deaths per year for most of the diseases. Pasteurellosis was reported to cause the most deaths for all ruminants and very similar mortalities were reported for Anthrax and FMD in all ruminants; up to 3,000 deaths were due to FMD (Figures 11A and 11C). There were also similarities between states for all ruminants, with the most deaths in Amhara and Oromia (Figures 11B and 11D).

## 4 Discussion

This study aimed to assess the available data for mortality and reproductive losses, and their causes in cattle and small ruminants in states of Ethiopia. Predictive models were created to analyse three datasets. Cattle were analysed separately to sheep and goats, which were combined due to their similarities. Deaths and abortions were analysed separately.

Across states and years, most herds reported no deaths, with only rare occasions of large losses. In herds with increased mortality the cause of death in both cattle and SR varied by State, Years and Age groups and Production System. In cattle herds, mortality in young stock was more commonly associated with respiratory and gastrointestinal disease. In SR herds, mortality in both young stock and adults was commonly associated with respiratory and gastrointestinal diseases. In both cattle and SR herds, pastoralists reported the highest mortality. Wide variation in the predicted number of deaths in both cattle and SR was seen across agroecological zones; in 2011 in Oromia, most cattle deaths occurred in tropic cool semi-arid areas, compared to in 2013 when most occurred in tropic cool sub-humid areas. There was similar variation for SR deaths across agroecological zones but no clear trend in the association of mortality with either humidity or temperature.

Unsurprisingly, number and reasons for abortions in both cattle and SR were associated with disease, eco-system, time and place. Given relatively small herd sizes, we predicted zero abortion for most herds. Increased numbers of abortions were predicted for cattle herds in the lowlands of Oromia and Somali and predicted for SR in the lowlands of Amhara. Where the cause of the abortion was known, abortions in both cattle and SR were most likely caused by brucellosis while in cattle abortions were appreciably also caused by FMD.

The data can be considered ‘noisy’, with such a high degree of variation that interpretation of them is challenging. It is necessary to question why the data are noisy; are the underlying data unreliable, or is natural/ environmental variation the cause of the variability?

With regards to the reliability of the data, there were indeed some areas of concern. From the Farmer Surveys, it was reported that sometimes respondents were not able to provide reliable information about the numbers of their animals and sometimes provided exaggerated mortality figures, not matching the actual herd size. Key informant discussion suggested that disease outbreak reports were not carefully or regularly recorded and recommended the expansion of reliable diagnostic tests, particularly field tests. Additionally, improvements in capacity building and practical training were recommended, in order to improve the quality and quantity of OIE disease reporting. The overall impression of the diagnostic laboratory data supplied was that it lacked detail and did not appear comprehensive, rather providing a reflection of the general causes of mortality and reproduction losses. Several key informants recommended the strengthening of regional laboratories by fulfilling equipment needs and necessary funding, and suggesting that laboratories needed to strengthen their diagnostic capacity. A recent study reported that a lack of diagnostic laboratory resources, combined with the presence of most potential infectious and non-infectious causes of abortion made diagnosis particularly challenging in Ethiopia (Gelalcha et al., 2021).

The greater mean number of deaths predicted for cattle and SR herds in Oromia are likely caused by the greater proportion of pastoralist herds with higher number of animals in that area. Other likely sources of natural variation include different conditions for different years, such as droughts and epidemics (Alemayehu and Fantahun, 2012; Megersa et al., 2014). That there was evidence of spatiotemporal variation of risk factors and causes of mortality is not surprising, as seasonal and environmental variation affects grazing availability and ensuing movement and mixing of animals (Devereux, 2006; Getachew et al., 2010; Gelalcha et al., 2021) which can then affect disease transmission. A recent systematic review observed a scarcity of disease-associated mortality data for Ethiopia, with only 14 studies identified (Tsouloufi et al., 2022, submitted). In our study’s semis-tructured interviews, a key informant reported drought as one of the major causes for mortality and reproductive losses in cattle and small ruminants, with peak mortality with the first rains following drought. Droughts occur frequently in Ethiopia, causing severe damage and harm. Previously, droughts were reported to occur every 8-10 years for the whole country but recently it has been observed that the frequency of drought has shortened, with severe drought happening three years in succession, which was the situation in 2011 and 2015-2017 (USAID, 2019). Conversely, the relative lack of variability in the DOVAR dataset was surprising, and as this data collection only captured disease-related events, it is difficult to interpret.

While infectious causes are regarded as predominant in ruminant abortions (Sebastiani et al., 2018), environmental factors are known risk factors (Menzies, 2011), as are management systems and husbandry factors (Mekonnen et al., 2010). As husbandry systems vary across agroecological zones (Fournié et al., 2018) disease prevalence varies widely across agroecological zone, as shown by Welay *et al.* (2018). Therefore, seasonality, agroecology and production systems all significantly affect abortions (Alemayehu et al., 2021). A recent review of the causes for small ruminant abortion in Ethiopia reported several infectious diseases and non-infectious conditions, and highlighted evidence gaps on disease dynamics (Gojam and Tulu, 2020). It also identified the major risk factors for small ruminant abortion, including age, parity, genetics, geographical and environmental factors, and herd size in the varying production systems. In order to establish the source of the variability, it would be necessary to compare the data with unbiased data, something that was not available. However, as multiple data sources were assessed and similar heterogeneity was observed in those analysed, it is likely that the observed heterogeneity is real, but possibly confounded with, and compounded by data collection artefacts.

It was notable that the addition of some explanatory factors caused models to fail to converge, however, once other variables had been added it was sometimes possible to include them. The reported models should be viewed as being the optimum models to describe the data, given the datasets provided, and it cannot be guaranteed that replications of the datasets would result in the same models being selected.

Using the models to examine predictive outputs i.e. how many deaths or abortions we would expect from a similar set of herds across the various explanatory factors shows that in nearly all cases zero deaths or abortions would be observed. There are, however, occasional herds where relatively large numbers of deaths or abortions might occur; in the farmer survey, for example, up to 90 deaths were predicted in a single epidemiological unit. The data did not provide the evidence required to explore whether this variability is consistent across time or not.

As stated, the Farmers Surveys and LSMS analyses indicated that losses, from both deaths and abortions, in domesticated ruminants in Ethiopia demonstrate a large amount of variability, both geographically and across time. They represent losses from a number of diseases, and the major diseases recorded vary across both time and space. It is likely that the country-based mean death rate varies widely across different areas, management systems and from year to year. The fitted models assist our understanding of patterns of mortality, however, the degree of variation observed, particularly across time, suggests that these models are only really useful as a *descriptive* tool, and are unlikely to be useful *predictively*. The model predictions provide some insight into why mortality rates are highly variable and therefore unlikely to be a useful measure of production or change in health status. As most herds had no deaths or abortions, with a few herds having relatively high mortality, a single metric is very hard to interpret. This was reflected in our modelling strategies, and in particular in the choice of hurdle models and models allowing for zero-inflation and excess heterogeneity.

Summaries would be best based on data that are more likely to be stable, i.e. less likely to change due to random influences. Furthermore, for farmers to keep reliable records of outcomes, they should be encouraged to record outcomes that are of interest to the farmer. While deaths and abortions ought to be of interest to farmers, their rarity and high variability likely make them of less use to a researcher trying to improve productivity, and indeed they are only indirectly related to productivity as they represent a potential loss. Tsouloufi *et al.* recently established a systematic mapping protocol to review the available evidence on disease prevalence and associated mortality in ruminants in Ethiopia, in attempt to address the disparate data and to improve decision-making on disease control policies (Tsouloufi et al., 2020). A recent systematic review by Wong *et al.* (2021) observed that the greatest proportion of mortalities in cattle, sheep and goats was reported to be in those under the age of six months. Interestingly, this study did not observe increased mortality in young stock compared to adults, as young stock mortality is generally considered high in sub-Saharan Africa, which leads us to question farmer recall and whether they remember adult losses more than young stock. Wong *et al.* (2021) also highlighted that there was a high degree of variability in both causes and risk factors which meant that ‘mortality rate’ was an incomplete indicator, and as such, qualitative and quantitative data on mortality causes would improve the understanding and allow preventative actions. Information is required on the causes of mortality for decision-making, as well as the rates in different livestock species (Catley et al., 2014).

We would suggest, therefore, the investigation of alternative measures of productivity. For example, number of live births per dam per year, number of offspring raised to one year old per dam, or median or mean age of adults are likely to be more indicative and useful measures. The first relates to fertility, embryonic death, abortion and stillbirth; the second to mortality of young stock; the last to adult mortality. All three variates would be expected to be more stable over time than numbers of deaths or abortions, since all observational units (herds) contribute to the estimate, and not merely those with large losses. In addition, they are likely of more direct interest to farmers since they represent profit or productivity, and can be more easily equated to financial gain/loss.

The study had several limitations. The objectives were somewhat misguided in that they were initially to obtain numbers in order to inform interventions. None of the datasets provided could be reasonably expected to provide unbiased, accurate estimates of the death or abortion rate. The farmer survey was a convenience survey, using local experts to help define the sampling frame. Thus, the local experts might have chosen individuals known to have had problems, or not. The LSMS data comes from an unknown sampling frame. It is believed that this sampling frame is carefully chosen, but it is meant to represent a country’s economy, and is not necessarily representative of its agricultural sector or livestock population. Additionally, these data are collected retrospectively; all individuals are asked for their losses during the previous year, and hence are subject to recall bias. The DOVAR dataset, although potentially unbiased, requires a chain of events to occur before the data become available; the animal must first contract a notifiable disease; someone must recognize the disease, then decide to report it; the report must be passed through a reporting chain until it is collated into the country-wide statistic.

## Conclusion

The study contributes to the understanding of the patterns of mortality in domesticated ruminants in Ethiopia. The degree of variation observed, especially temporally, suggests that the predictive nature of the models is limited and that the estimated mortality rates are highly variable and therefore lack reliability as a metric of productivity. Unrecorded variables such as changeable seasons and environmental conditions likely affect the morbidity and mortality rates across space and time and thus the high variability observed is challenging to interpret. Nevertheless, whilst there are many limitations in the use of ‘noisy data’, there are also many opportunities; indeed, noisy data may be the best available, and can provide valuable insights otherwise untold of the situation in the field, especially in rural settings in LMICs, with limited resources. As such, this study should inform the choice of dataset for future metrics, such as young stock mortality, and the choice of statistical method used to calculate metrics that may better inform productivity and allow for suitable preventative actions and interventions to be implemented as required.

## Supporting information

Supplementary Farmer Surveys

## 5 Data Availability Statement

The datasets for this study can be found in the GitLab https://gitlab.bioss.ac.uk/giles/sebi-data.

## 6 Author Contributions

CV and AP designed the study. SH and VN carried out the field work. GT, DE, IM, FA and CS undertook analysis and interpretation. All authors wrote the manuscript.

## 7 Funding

This work was supported by the Bill and Melinda Gates Foundation [OPP1134229, to SEBI-Livestock]. The funders had no role in the study design, data collection and analyses, decision to publish, or preparation of the manuscript.

## 8 Conflict of Interest

The authors declare that the research was conducted in the absence of any commercial or financial relationships that could be construed as a potential conflict of interest.

## 9 Acknowledgments

The authors wish to thank all livestock keepers for their participation. Thanks are also given to Dr Alemayehu Mekonnen Anbessie, former CVO/OIE delegate of Ethiopia, current Senior Advisor to office of the Minister for Livestock and Fisheries Resources Development Sector, MOA, Ethiopia, and to the Ethiopian Ministry of Agriculture and Livestock. Gratitude is also extended to Federal Livestock and Fisheries Development staff, particularly Misrak Mekonnen, Thomas Cherinet and Yismashewa Wegayehu.

## 10.1 Supplementary Material

Farmer surveys for cattle and small ruminants are shown in Supplementary Material.

## Footnotes

1 Since the surveys were conducted, a tenth regional state has been created

## Notes

### Competing Interest Statement

The authors have declared no competing interest.

