## Supplementary Farmer Surveys for "Patterns of mortality in domesticated ruminants in Ethiopia"

Supplementary Material

Farmer Survey for cattle owners

**Surveillance of major infectious and reproductive diseases of cattle**

DEMOGRAPHIC DATA

**1. Name (optional) …………………………………………	phone number ………………………………….**

**2. (Location) State……………………. Local Government Area……………………………………………**

**3. Age ………………………………. Educational level …………………………………………………………….**

**4. How long have you been keeping ruminants? …………………………………………………………….**

**5. What kind of animals do you keep (Please Tick as appropriate) Beef Cattle 🞏 Dairy Cattle 🞏**

MORTALITY AND REPRODUCTIVE LOSSES

| **Serial** | **Questions** | **Bulls** | **Lactating cow** | **Dry cows** | **Heifers** | **Steers** | **Calves** | **Oxon** |
| --- | --- | --- | --- | --- | --- | --- | --- | --- |
| **6** | **Average herd composition last year** |  |  |  |  |  |  |  |

| **Serial** | **Questions** | **Bulls** | **Lactating cow** | **Dry cows** | **Heifers** | **Steers** | **Calves** | **Oxon** |
| --- | --- | --- | --- | --- | --- | --- | --- | --- |
| **7** | **Animals dead last year** |  |  |  |  |  |  |  |
|  | **Tryps** |  |  |  |  |  |  |  |
|  | **FMD** |  |  |  |  |  |  |  |
|  | **LSD** |  |  |  |  |  |  |  |
|  | **Infectious respiratory diseases:**  ***CBPP**  ***Pasteurellosis**  ***Other infectious respiratory diseases** |  |  |  |  |  |  |  |
|  | **Injuries** |  |  |  |  |  |  |  |
|  | **Nutrition (insufficient food/water)** |  |  |  |  |  |  |  |
|  | **GI parasites** |  |  |  |  |  |  |  |
|  | **Other respiratory** |  |  |  |  |  |  |  |
|  | **Other digestive** |  |  |  |  |  |  |  |
|  | **Other reproductive** |  |  |  |  |  |  |  |
|  | **Sudden death** |  |  |  |  |  |  |  |
|  | **Regular slaughter** |  |  |  |  |  |  |  |
|  | **Other - unspecific** |  |  |  |  |  |  |  |

| **Serial** | **Questions** | **Bulls** | **Lactating cow** | **Dry cows** | **Heifers** | **Steers** | **Calves** | **Oxon** |
| --- | --- | --- | --- | --- | --- | --- | --- | --- |
| **8** | **Number of abortions last year** |  |  |  |  |  |  |  |
|  | **Clinical signs compatible with Brucellosis** |  |  |  |  |  |  |  |
|  | **Clinical signs compatible with FMD** |  |  |  |  |  |  |  |
|  | **Clinical signs compatible to other diseases** |  |  |  |  |  |  |  |
|  | **Linked to heat stress** |  |  |  |  |  |  |  |
|  | **Linked to injuries and other causes** |  |  |  |  |  |  |  |
|  | **Linked to poor nutrition** |  |  |  |  |  |  |  |
|  | **Early abortions (< 5 months)** |  |  |  |  |  |  |  |
|  | **Late abortions (> 5 months)** |  |  |  |  |  |  |  |

| **Serial** | **Questions** | **Bulls** | **Lactating cow** | **Dry cows** | **Heifers** | **Steers** | **Calves** | **Oxon** |
| --- | --- | --- | --- | --- | --- | --- | --- | --- |
| **9** | **Animals sold because illness last year** |  |  |  |  |  |  |  |
|  | **Tryps** |  |  |  |  |  |  |  |
|  | **LSD** |  |  |  |  |  |  |  |
|  | **FMD** |  |  |  |  |  |  |  |
|  | **Infectious respiratory diseases:**  ***CBPP**  ***Pasteurellosis**  ***Other infectious respiratory diseases** |  |  |  |  |  |  |  |
|  | **Injuries** |  |  |  |  |  |  |  |
|  | **Nutrition (insufficient food/water)** |  |  |  |  |  |  |  |
|  | **GI parasites** |  |  |  |  |  |  |  |
|  | **Other respiratory** |  |  |  |  |  |  |  |
|  | **Other digestive** |  |  |  |  |  |  |  |
|  | **Other reproductive** |  |  |  |  |  |  |  |
|  | **Other - unspecific** |  |  |  |  |  |  |  |

**Thank You for Your Participation**

**Farmer Survey for sheep and goat owners**

**Surveillance of major infectious and reproductive diseases of small ruminants**

DEMOGRAPHIC DATA

**1. Name (optional) …………………………………………	phone number ………………………………….**

**2. (Location) State……………………. Local Government Area……………………………………………**

**3. Age ………………………………. Educational level …………………………………………………………….**

**4. How long have you been keeping ruminants? …………………………………………………………….**

**5.1 What kind of animals do you keep (Please Tick as appropriate) Sheep 🞏	Goat 🞏**

| **Serial** | **Question** | **Sheep** | **Goat** |
| --- | --- | --- | --- |
| **5.2** | **Total population last year** |  |  |

MORTALITY AND REPRODUCTIVE LOSSES

| **Serial** | **Question** | **< 3m male** | **< 3m female** | **3m - 1 yr male** | **3m - 1 yr female** | **> 1 yr male** | **> 1 yr female** |
| --- | --- | --- | --- | --- | --- | --- | --- |
| **6** | **Average herd composition last year** |  |  |  |  |  |  |

| **Serial** | **Questions** | **< 3m male** | **< 3m female** | **3m - 1 yr male** | **3m - 1 yr female** | **> 1 yr male** | **> 1 yr female** |
| --- | --- | --- | --- | --- | --- | --- | --- |
| **7** | **Animals dead last year** |  |  |  |  |  |  |
|  | **Tryps** |  |  |  |  |  |  |
|  | **Orf** |  |  |  |  |  |  |
|  | **FMD** |  |  |  |  |  |  |
|  | **Infectious respiratory diseases:**  ***CCPP** |  |  |  |  |  |  |
|  | ***PPR** |  |  |  |  |  |  |
|  | ***Pasteurellosis** |  |  |  |  |  |  |
|  | ***Other infectious respiratory diseases** |  |  |  |  |  |  |
|  | **Injuries** |  |  |  |  |  |  |
|  | **Foot rot** |  |  |  |  |  |  |
|  | **Nutrition (insufficient food/water)** |  |  |  |  |  |  |
|  | **GI parasites** |  |  |  |  |  |  |
|  | **Other respiratory** |  |  |  |  |  |  |
|  | **Other digestive** |  |  |  |  |  |  |
|  | **Other reproductive** |  |  |  |  |  |  |
|  | **Sudden death** |  |  |  |  |  |  |
|  | **Other - unspecific** |  |  |  |  |  |  |

| **Serial** | **Questions** | **< 3m male** | **< 3m female** | **3m - 1 yr male** | **3m - 1 yr female** | **> 1 yr male** | **> 1 yr female** |
| --- | --- | --- | --- | --- | --- | --- | --- |
| **8** | **Number of abortions last year** |  |  |  |  |  |  |
|  | **Clinical signs compatible with Brucellosis** |  |  |  |  |  |  |
|  | **Clinical signs compatible with FMD** |  |  |  |  |  |  |
|  | **Clinical signs compatible with PPR** |  |  |  |  |  |  |
|  | **Clinical signs compatible to other diseases** |  |  |  |  |  |  |
|  | **Linked to heat stress** |  |  |  |  |  |  |
|  | **Linked to injuries and other causes** |  |  |  |  |  |  |
|  | **Linked to poor nutrition** |  |  |  |  |  |  |

| **Serial** | **Questions** | **< 3m male** | **< 3m female** | **3m - 1 yr male** | **3m - 1 yr female** | **> 1 yr male** | **> 1 yr female** |
| --- | --- | --- | --- | --- | --- | --- | --- |
| **9** | **Animals sold because illness last year** |  |  |  |  |  |  |
|  | **Tryps** |  |  |  |  |  |  |
|  | **Orf** |  |  |  |  |  |  |
|  | **FMD** |  |  |  |  |  |  |
|  | **Infectious respiratory diseases:**  ***CCPP** |  |  |  |  |  |  |
|  | ***PPR** |  |  |  |  |  |  |
|  | ***Pasteurellosis** |  |  |  |  |  |  |
|  | ***Other infectious respiratory diseases** |  |  |  |  |  |  |
|  | **Injuries** |  |  |  |  |  |  |
|  | **Foot rot** |  |  |  |  |  |  |
|  | **Nutrition (insufficient food/water)** |  |  |  |  |  |  |
|  | **GI parasites** |  |  |  |  |  |  |
|  | **Mange** |  |  |  |  |  |  |
|  | **Other respiratory** |  |  |  |  |  |  |
|  | **Other digestive** |  |  |  |  |  |  |
|  | **Other reproductive** |  |  |  |  |  |  |
|  | **Other - unspecific** |  |  |  |  |  |  |

**Thank You for Your Participation**
